# Mechano-activated synovial cells instigate OA-like disease progression in the absence of joint injury

**DOI:** 10.64898/2026.10.06.756770

**Authors:** Elizabeth R. Bernstein, Georgios Kotsaris, Alexandra A. Silverman, Tsegaw Bahlibi, Brett Croen, Rachel Flaugh, Carla Scanzello, Liane Miller, Joel D. Boerckel, Nathaniel A. Dyment, Robert L. Mauck

**Affiliations:** Department of Bioengineering, University of Pennsylvania, Philadelphia, PA, United States; Department of Orthopaedic Surgery, University of Pennsylvania, Philadelphia, PA, United States; CReATE Motion Center, CMC VA Medical Center, Philadelphia, PA, United States; Department of Medicine, Division of Rheumatology, University of Pennsylvania, Philadelphia, PA, United States

**Keywords:** Synovial mechanobiology, Osteoarthritis, YAP/TAZ signaling

## Abstract

While altered tissue mechanics and cellular mechanoactivation accompany osteoarthritis, whether dysregulated mechanotransduction can initiate joint disease remains unclear. Here, we used intra-articular delivery of cell-penetrating TAT-Cre recombinase to locally perturb the actomyosin–Hippo mechanotransduction axis in mice. Deletion of *Lats1/2*, the negative regulators of the mechanosensitive transcriptional coactivators YAP and TAZ, induced sustained joint hyperalgesia, altered gait, synovial hyperplasia and fibrosis, vascular remodeling, and articular cartilage degeneration in the absence of a destabilizing injury. Single-cell transcriptomics revealed the emergence of *Cthrc1*^+^ matrix-remodeling fibroblasts and accumulation of *Mrc1^+^Trem2^+^*macrophages during this pathological process. Histological analyses showed persistent contractile stromal remodeling and an early macrophage response that partially subsided over four weeks. Conversely, local deletion of *Myh9/10*, encoding the force-generating heavy chains of non-muscle myosin II, reduced hyperalgesia, synovial lining hyperplasia, αSMA^+^ cell accumulation, and macrophage infiltration during post-traumatic osteoarthritis (PTOA) pathogenesis. However, this intervention did not detectably reduce cartilage degeneration at eight weeks. These findings establish synovial mechanotransduction as a causal driver of joint disease via immune–stromal remodeling and pain development. In addition, partial suppression of contractility separates symptomatic from structural outcomes in PTOA. Synovial mechanotransduction therefore represents a target for modulating pathological stromal–immune remodeling and joint pain.

## INTRODUCTION

The mechanical environment is a fundamental regulator of cell behavior, shaping phenotype and function across nearly every tissue in the body^1^. Cells sense and respond to these cues in large part through their cytoskeleton, which couples extracellular matrix engagement to intracellular force generation and downstream transcriptional responses^2^. This mechanotransduction process is essential for tissue formation, growth, and homeostasis^3,4^. Importantly, cells often adopt a mechanoactivated or mechano-inflammatory state during specific phases of disease pathogenesis^5,6^. As an example, fibrotic remodeling is accompanied by extracellular-matrix accumulation and tissue stiffening, which reinforce cellular contractility, proliferation, and matrix production through mechanoresponsive signaling^7^. Thus, cellular mechanoactivation and matrix remodeling may form a feed-forward relationship during disease progression. That said, it is not clear whether mechano-activation itself can independently cause disease or whether it is merely a consequence of disease and/or a facilitator of disease pathogenesis and severity.

A central regulator of mechanoactivation across cell types is the actomyosin–Hippo pathway^8–10^. Non-muscle myosin II (NM-II) engages with filamentous actin to generate intracellular contractile forces, while the Hippo pathway kinases LATS1 and LATS2 restrain the mechanosensitive transcriptional co-activators YAP and TAZ^8,11^. Increased cytoskeletal tension suppresses LATS1/2 activity, permitting YAP/TAZ nuclear accumulation and induction of mechanoresponsive transcriptional programs^12^. In fibroblasts, sustained YAP/TAZ activity promotes contractility, extracellular matrix production, and myofibroblast differentiation while, in macrophages, it regulates inflammatory activation and migration^13–15^. Thus, the actomyosin– Hippo pathway provides a shared mechanistic framework through which stromal and immune cells respond to altered tissue mechanics.

This mechanotransduction axis is particularly relevant to osteoarthritis (OA), a debilitating whole-joint disease affecting hundreds of millions of people worldwide for which no disease-modifying therapies currently exist^16,17^. Although cartilage degeneration has historically dominated the study of OA, synovial inflammation, hyperplasia, neovascularization, and fibrosis are increasingly recognized as important contributors to disease progression and pain^18^. Notably, progressive extracellular matrix deposition, cell proliferation, and overall tissue hypertrophy are hallmarks of fibrotic synovium that transform this normally compliant tissue into a stiff and mechanically dysregulated microenvironment, potentially establishing a feed-forward loop in which tissue stiffening promotes cellular mechanoactivation, which in turn drives further inflammation and matrix deposition^19–21^. Consistent with this model, OA is associated with progressive stiffening of the synovial microenvironment and mechanoactivation of fibroblast and macrophage populations^20,21^. However, these observations remain correlative and do not establish whether dysregulated synovial cell mechanotransduction is sufficient to initiate joint pathology. Specifically, it remains unknown whether sustained activation of synovial mechanotransduction is sufficient to initiate joint pathology in the absence of injury, how such perturbation reshapes fibroblast and macrophage states within the synovium, and whether suppressing cytoskeletal mechanotransduction can attenuate disease progression following joint injury. Resolving these questions would establish whether altered tissue mechanics are simply a consequence of pathological remodeling or whether dysregulated mechanotransduction can itself initiate and drive disease pathogenesis.

Here, we addressed these questions by locally manipulating complementary nodes of the actomyosin–YAP/TAZ mechanotransduction axis using intra-articular delivery of the cell-penetrating TAT-Cre recombinase^22^. We first deleted *Lats1/2* in the uninjured joint to induce sustained YAP/TAZ activation and determine whether aberrant mechanotransduction is sufficient to initiate joint pathology. Local *Lats1/2* deletion induced pain, synovial hyperplasia, fibrosis, vascular remodeling, and progressive joint degeneration in the absence of a destabilizing injury. Single-cell transcriptomic profiling revealed coordinated remodeling of both fibroblast and macrophage populations, linking sustained mechanoactivation to altered stromal and immune cell states, consistent with mechano-inflammation. Conversely, to determine whether suppressing cytoskeletal mechanotransduction could mitigate injury-induced disease, we deleted NM-II to reduce actomyosin contractility, which attenuated synovial pathology and pain following destabilization of the medial meniscus. Together, these findings identify synovial mechanotransduction and mechano-inflammation as a causal regulator of pathological immune– stromal remodeling and demonstrate that opposing perturbations of the actomyosin–YAP/TAZ axis can profoundly alter joint pathology.

## RESULTS

### Intra-articular *Lats1/2* ablation drives joint pathology in the absence of injury

TAT-mediated protein transduction is a novel method for delivering recombinant Cre to cells and tissues that allows spatially- and temporally-controlled genetic modulation^22^. As we aimed to use this approach to achieve genetic ablation in the synovium, we first confirmed that TAT-Cre delivery to synovial fibroblasts in culture resulted in efficient recombination (recombination rate = 79.9%) (**Supplementary Fig. 1A,B**). Then, we employed the TAT-Cre system to assess the extent of local genetic recombination induced by intra-articular TAT-Cre delivery in tdTomato Ai9 reporter mice. Following intra-articular injection, PBS-injected knees showed minimal tdTomato fluorescence, whereas TAT-Cre-injected knees exhibited robust tdTomato signal throughout the synovium (**Fig. 1A**). Minimal, superficial tdTomato signal was detected in the articular cartilage and in the anterior and posterior cruciate ligaments, indicating that recombination was largely targeted to the synovial compartment. Flow cytometry detected tdTomato in 22.4% of CD45*^+^* immune cells and 54.8% of CD45^-^ stromal cells after TAT-Cre delivery (**Supplementary Fig. 1C,D**), with only 1.2% and11.3% tdTomato^+^ cells detected within these gates in PBS-injected controls. Thus, intra-articular TAT-Cre enabled local, synovium-biased recombination in the absence of surgical injury.

**Fig. 1:**
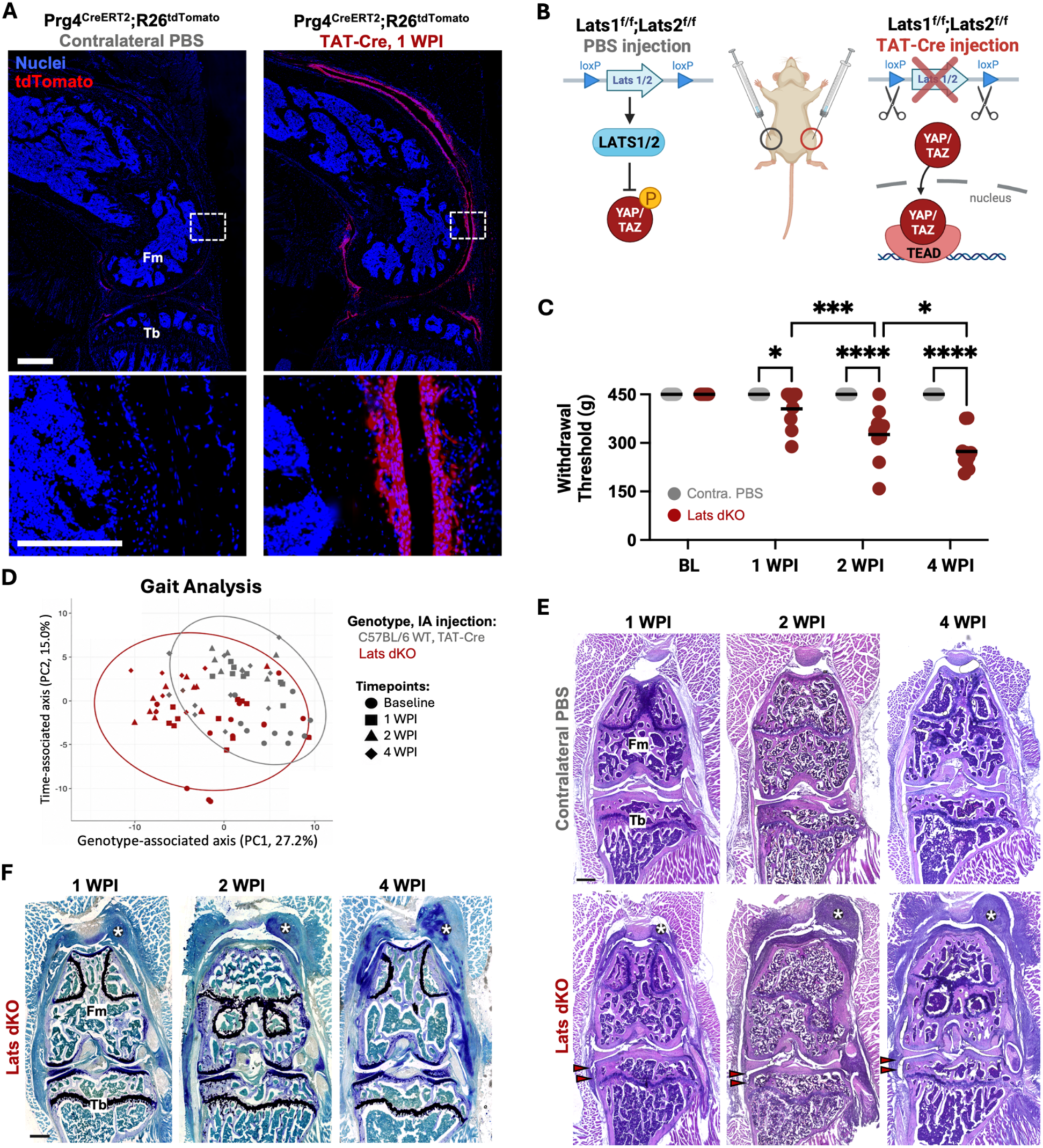
Local *Lats1/2* deletion induces pain, gait alterations, and progressive joint remodeling in the absence of injury. **(A)** Representative fluorescence images of PBS- and TAT-Cre-injected tdTomato Ai9 reporter knees showing tdTomato (red) and nuclei (blue), Scale = 500μm; lower panels show the boxed regions at higher magnification (Scale = 200μm). **(B)** Schematic of intra-articular PBS or TAT-Cre administration in Lats floxed mice and consequent regulation of YAP/TAZ activity. **(C)** Joint withdrawal thresholds at baseline and 1, 2, and 4 WPI in PBS- and TAT-Cre-injected Lats dKO knees. **(D)** Principal component analysis of gait parameters from TAT-Cre-injected wild-type and Lats dKO mice at baseline and 1, 2, and 4 WPI. PC1 and PC2 account for 27.2% and 15.0% of the variance, respectively; colors indicate genotype and symbols indicate time point. **(E)** Representative H&E-stained whole-joint sections from PBS- and TAT-Cre-injected Lats dKO knees at 1, 2, and 4 WPI. * indicates peri-articular fibrous/chondroid masses. **(F)** Representative Toluidine blue staining of TAT-Cre-injected Lats dKO knees at 1, 2, and 4 WPI. Scale = 500 μm. Fm, femur; Tb, tibia. *p<0.05, ***p<0.001, ****p<0.0001.

To determine whether mechanoactivation of the synovium could initiate joint disease in the absence of injury, we targeted the Hippo regulators LATS1/2, whose function is to restrain mechanosensitive YAP/TAZ transcriptional activity^23,24^. This was accomplished via two repeated intraarticular injections of TAT-Cre (separated by 24hrs) to the knees of *Lats1*^f/f^;*Lats2*^f/f^ mice (hereafter, Lats dKO). Contralateral knees received PBS as internal controls (hereafter, contra PBS) (**Fig. 1B**). Additional wild-type mice were injected with TAT-Cre as controls for the effects of TAT-Cre. Pain-related behavior, gait analysis, and joint histopathology were evaluated at 1, 2, and 4 weeks post-injection (WPI). TAT-Cre-injected knees of Lats dKO mice exhibited lower withdrawal thresholds than contralateral PBS-injected knees at each time point (**Fig. 1C**), indicating sustained joint hyperalgesia. TAT-Cre did not alter withdrawal thresholds in wild-type mice, indicating that these findings were due to *Lats1/2* deletion rather than TAT-Cre exposure.

Knee hyperalgesia is often accompanied by changes in limb function and behavior^25–27^. To determine whether hyperalgesia was accompanied by altered limb function, we evaluated gait at baseline and at 1, 2, and 4 WPI. Principal component analysis partially separated Lats dKO and wild-type mice along PC1, which accounted for 27.2% of the total variance, while PC2 accounted for an additional 15.0% of the variance and reflected differences in assessment timepoint (**Fig. 1D**). These data indicate that local *Lats1/2* deletion produced a distinct, although heterogeneous, gait profile that evolved over time.

TAT-Cre-injected Lats dKO knees displayed marked synovial hypercellularity and thickening as early as 1 WPI, which progressed through 4 WPI (**Fig. 1E**). In addition, fibrotic nodules that extended from the joint capsule and into the periarticular muscle regions were frequently observed. Toluidine blue staining revealed chondroid matrix within these periarticular lesions, consistent with ectopic cartilage formation (**Fig. 1F**). Together, these findings show that local *Lats1/2* deletion is sufficient to induce pain, functional impairment, and progressive multi-tissue joint remodeling in the absence of a destabilizing injury.

### Intra-articular *Lats1/2* ablation drives synovial fibrosis, vascular remodeling, and articular cartilage degeneration in the absence of injury

Synovial hyperplasia and fibrosis are established features of osteoarthritis and can alter the mechanical environment experienced by resident cells^21,28,29^. To define the tissue changes accompanying the pain and gait phenotypes, we examined the synovium at higher resolution. At 2 WPI, TAT-Cre-injected Lats dKO knees exhibited pronounced intimal lining hyperplasia, increased sublining cellularity, and fibrosis in the synovial gutters (**Fig. 2A**). Semiquantitative scoring showed that each feature was already elevated at 1 WPI and remained elevated through 4 WPI (**Fig. 2C, Supplementary Fig. 2**), demonstrating rapid and persistent synovial remodeling.

**Fig. 2:**
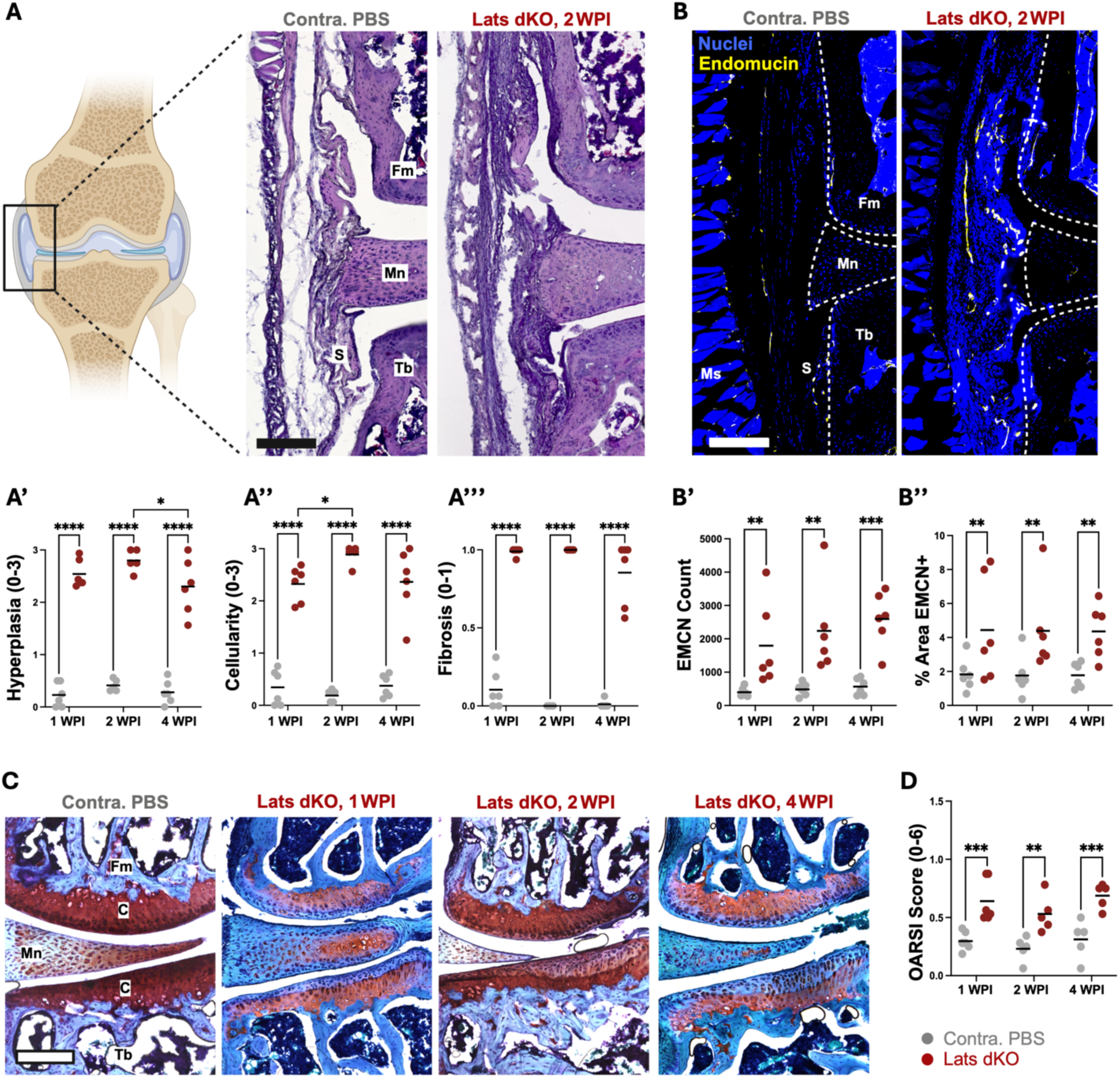
Intra-articular *Lats1/2* ablation drives synovial remodeling and cartilage degeneration in the absence of injury. **(A)** Representative H&E-stained sections (and scoring) of the synovial gutter of PBS- and TAT-Cre-injected Lats dKO knees at 2 WPI. **(A’-A’’’)** Semiquantitative scores for intimal hyperplasia, sublining cellularity, and fibrosis at 1, 2, and 4 WPI. **(B)** Representative fluorescence images (and quantification) of EMCN+ vascular structures in the synovial and parapatellar regions at 2 WPI. Nuclei are in blue and EMCN is in yellow. **(B’-B’’)** EMCN+ vessel count and EMCN+ percentage area at 1, 2, and 4 WPI. **(C)** Representative Safranin O/Fast green stained articular cartilage from PBS-injected control and TAT-Cre-injected Lats dKO knees at 1, 2, and 4 WPI. **(D)** OARSI scores at 1, 2, and 4 WPI. Scale = 200μm. C, cartilage; Fm, femur; Mn, meniscus; Ms, muscle; S, synovium; Tb, tibia. *p<0.05, **p<0.01, ***p<0.001, ****p<0.0001.

Synovial neovascularization is a characteristic feature of injury-associated synovitis and osteoarthritis and may facilitate immune-cell recruitment into the joint^30^. We therefore evaluated vascular remodeling by staining for the endothelial marker endomucin (EMCN). Representative images at 2 WPI showed increased vascular structures within the synovial gutters of TAT-Cre-injected Lats dKO knees (**Fig. 2B**). Quantification revealed a consistent 2-fold induction in EMCN^+^ vessel number and EMCN+ area at 1, 2, and 4 WPI, relative to PBS-injected controls (**Fig. 2D**).

As synovitis can precede cartilage lesions and is associated with subsequent cartilage loss during osteoarthritis progression^31–33^, we next asked whether the synovial remodeling induced by local *Lats1/2* deletion was accompanied by cartilage degeneration. Safranin O/Fast Green staining revealed preserved proteoglycan staining and surface architecture in PBS-injected knees, whereas Lats dKO knees displayed reduced Safranin O staining, surface irregularities, and elevated OARSI scores beginning at 1 WPI, and persisting through 4 WPI (Fig. 2e-f). These changes occurred despite minimal tdTomato labeling within the articular cartilage, supporting the interpretation that cartilage degeneration developed secondary to local recombination and remodeling within the synovial environment. In sum, local *Lats1/2* deletion in the absence of injury initiated persistent synovial fibrosis and vascular remodeling, together with articular cartilage degeneration that reached levels comparable to that seen with a destabilizing injury.

### *Lats1/2* deletion reshapes the immune and stromal cell landscape of the joint

Since *Lats1/2* deletion produced pathology across multiple joint and periarticular tissues, we developed a whole-joint dissociation and single-cell RNA-sequencing workflow to capture the broader cellular response. We applied this approach to whole knee joints collected 2 weeks after PBS or TAT-Cre injection. Unsupervised clustering of the combined dataset identified 16 populations spanning immune and non-immune compartments, annotated using canonical marker expression (**Supplementary Fig. 3A**). Most populations were represented in both conditions, with the Lats dKO joints yielding a modestly larger number of recovered cells (**Supplementary Fig. 3B**). We analyzed the CD45*^+^* and CD45^−^ compartments separately to resolve condition-associated changes within the immune and stromal populations.

Subclustering of CD45*^+^* cells identified 14 immune populations (**Fig. 3A**). Neutrophils and inflammatory monocytes did not show any expansion at this time point. In contrast, a macrophage population expressing *Mrc1* (encoding CD206), *Trem2*, *Cd163*, *C1qa*, *Apoe*, and *Folr2* was markedly increased in Lats dKO joints, but was nearly absent from PBS-injected controls (**Fig. 3B**, **Supplementary Fig. 3C**). In a non-surgical model of ACL rupture, *Mrc1*^+^*Trem*2^+^ macrophages that accumulated after injury have been associated with protection from joint degeneration^34^. Hereafter, we refer to the population identified in our dataset as *Mrc1*^+^*Trem*2^+^ macrophages. These findings suggest that *Lats1/2* deletion favored the evolution of a distinct mechano-inflammatory macrophage state rather than sustaining a broad expansion of acute inflammatory myeloid populations at 2 WPI.

**Fig. 3:**
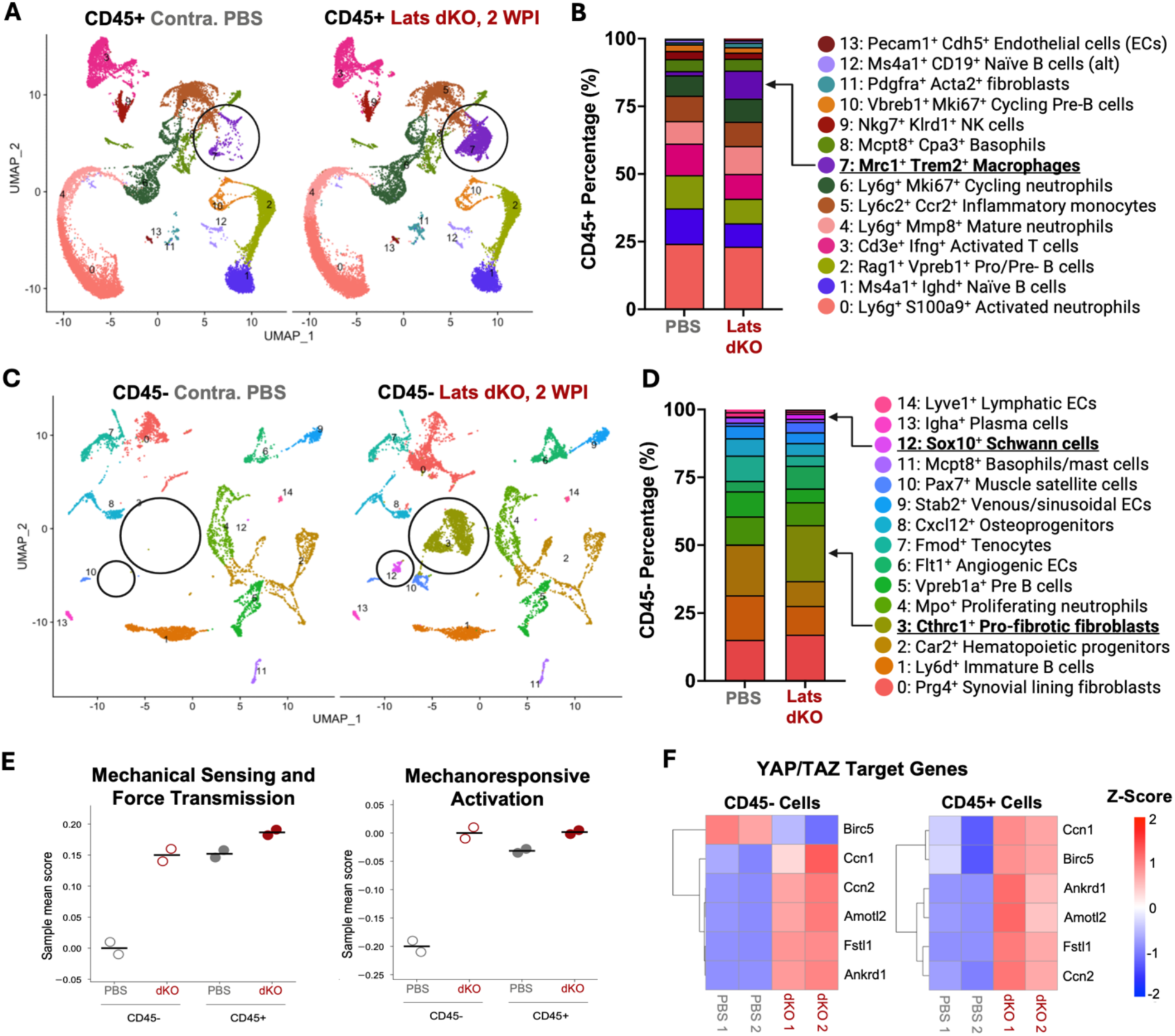
Local *Lats1/2* deletion reshapes joint immune and stromal cell landscape. **(A)** UMAP projections of CD45*^+^* cells from whole joints collected 2 WPI, colored by annotated population and displayed separately for PBS- and TAT-Cre-injected Lats dKO joints. Circle indicates the remodeling-associated macrophage cluster. **(B)** Relative composition of the CD45*^+^* compartment by condition. **(C)** UMAP projections of CD45^−^ cells, with circles indicating the *Cthrc1^+^*pro-fibrotic fibroblast and *Sox10^+^* Schwann cell populations. **(D)** Relative composition of the CD45^−^ compartment by condition. **(E)** Sample-level module scores for curated mechanosensing and mechanoactivation gene programs in CD45^−^ and CD45*^+^* cells. **(F)** Heatmaps showing scaled expression of select YAP/TAZ target genes in CD45^−^ and CD45*^+^* cells. Columns represent individual PBS or TAT-Cre samples and color denotes row-scaled expression.

Subclustering of CD45^−^ cells identified 15 stromal and vascular populations (**Fig. 3C**). Cluster 3, marked by *Cthrc1*, *Lox*, and *Pdpn*, was detected only in TAT-Cre-injected Lats dKO joints (**Fig. 3C,D**). *Cthrc1^+^* fibroblasts have been identified as matrix-remodeling populations in fibrotic tissues and in the arthritic pannus^35–37^, consistent with the synovial fibrosis observed histologically. The TAT-Cre dataset also showed greater representation of *Sox10^+^* Schwann cells and *Pax7^+^* muscle satellite cells (**Fig. 3D**), indicating that the response extended beyond the synovial fibroblast compartment and into periarticular neural and muscle-associated populations. These data indicate that local *Lats1/2* deletion induced a pathological fibroblast state while broadly altering the stromal and non-immune landscape of the joint.

We next tested whether these compositional changes were accompanied by increased mechanotransduction. Sample-level module scores for mechanical sensing and force transmission (*Piezo1, Trpv4, Itgb1, Tln1, Vcl,* and *Rock1/2*, among others) and mechanoresponsive activation (*Ccn1, Ccn2, Ankrd1, Amotl2,* and *Spp1*, among others) were higher in both the CD45⁻ and CD45⁺ compartments from TAT-Cre-injected joints (**Fig. 3E**). Consistent with LATS1/2 loss, CD45^−^ cells also showed coordinated increases in the YAP/TAZ target genes *Ccn1, Birc5, Cenpf, Ankrd1, Amotl2,* and *Fn1* (**Fig. 3F**). CD45^+^ cells exhibited a comparable response, including increased *Ccn1, Birc5, Ankrd1, Amotl2*, and *Fn1,* together with *Ccn2* (**Fig. 3F**). The transcriptional response was most pronounced in the stromal compartment, consistent with the prominent fibroproliferative phenotype observed histologically.

Together, these data demonstrate that local *Lats1/2* deletion reshaped the immune and stromal cell landscape, marked by the emergence of a *Cthrc1*⁺ matrix-remodeling fibroblast population and the accumulation of *Mrc1*^+^*Trem*2^+^ macrophages, alongside coordinated activation of mechanical sensing, force-transmission, and mechanoresponsive transcriptional programs.

### *Lats1/2* deletion produces sustained stromal and macrophage remodeling

To spatially validate the cellular changes identified by single-cell RNA sequencing, we stained joint sections collected at 1, 2, and 4 WPI for α-smooth muscle actin (αSMA), the macrophage marker F4/80, and EMCN (**Fig. 4A, Supplementary Figs. 5, 6**). Quantification was performed within the segmented pathological tissue region shown in **Fig. 4B**. αSMA-associated fluorescence increased ∼2-3-fold following *Lats1/2* deletion and remained elevated through 4 WPI (**Fig. 4C**), consistent with sustained expansion of contractile stromal populations. These αSMA*^+^* cells progressively extended into capsular and periarticular fibrotic regions. Because αSMA also labels vascular smooth muscle cells and pericytes, this signal should be interpreted as contractile stromal remodeling rather than a fibroblast-specific measurement.

**Fig. 4:**
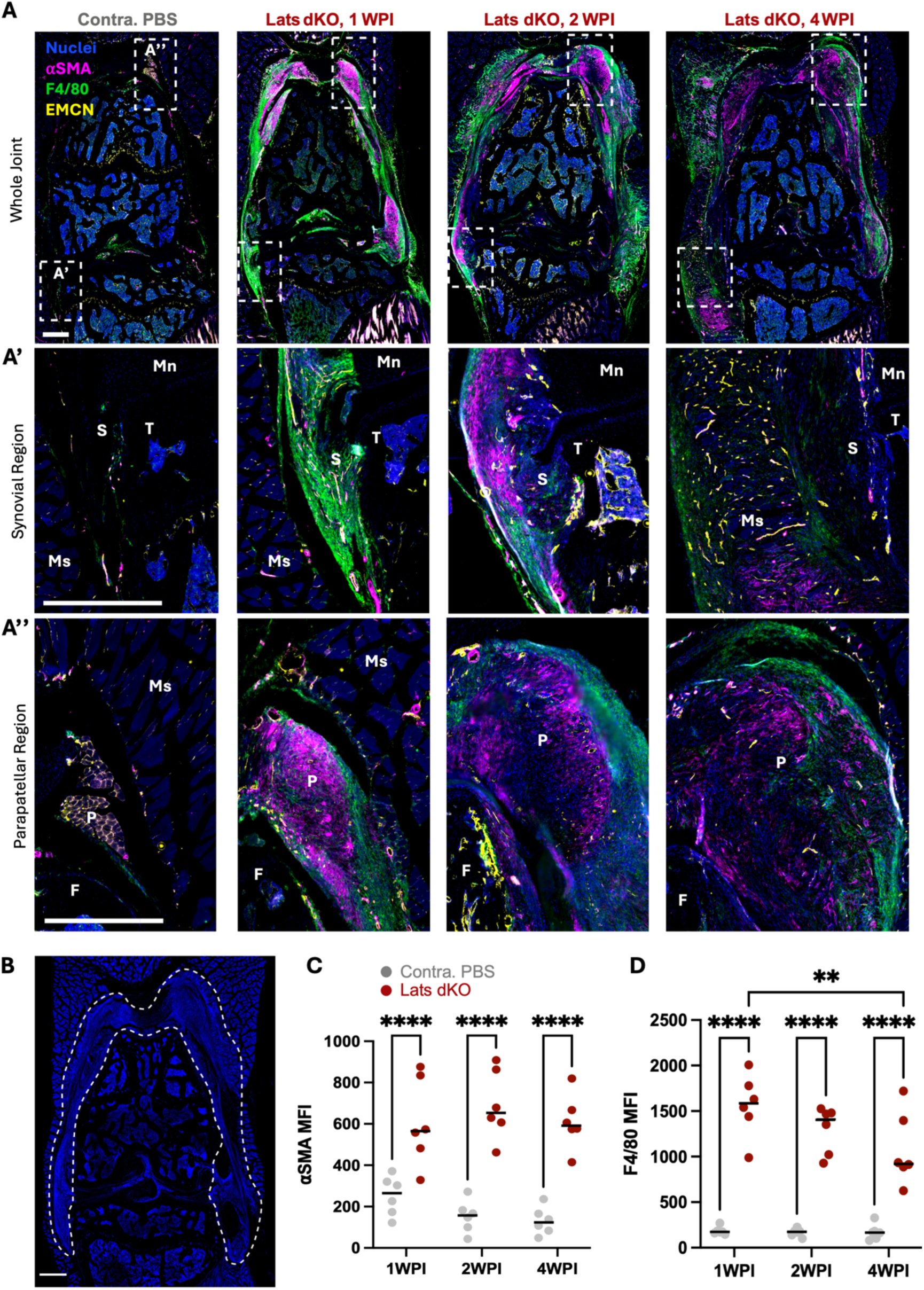
Local *Lats1/2* deletion results in sustained stromal and macrophage remodeling. **(A)** Representative immunofluorescence images of PBS-injected control knees and TAT-Cre-injected Lats dKO knees at 1, 2 and 4 WPI. Whole-joint views are shown above enlarged **(A’)** synovial and **(A’’)** parapatellar regions. Nuclei are in blue, αSMA is in magenta, F4/80 is in green and EMCN is in yellow. Boxes identify the enlarged regions. **(B)** Representative mask defining the pathological tissue region of interest (ROI) used for fluorescence quantification. Mean fluorescence intensity (MFI) of **(C)** αSMA and **(D)** F4/80 in the quantified ROI at 1, 2 and 4 WPI. Scale = 500 μm. C, capsule; F, femur; Mn, meniscus; Ms, muscle; P, patella; S, synovium; T, tibia. **p<0.01, ****p<0.0001.

Similar to αSMA, F4/80-associated fluorescence was also elevated at all time points compared to PBS controls, although this signal declined from a peak at 1 WPI by 4 WPI (**Fig. 4D**). Thus, mechano-inflammatory macrophage accumulation showed partial resolution but did not return to baseline within the four-week observation period. However, the macrophage response was substantially stronger than the αSMA response, with F4/80 having an 8.5-fold increase at 1 WPI, and a 7.8-fold increase at 2 WPI. Together with the scRNA-seq data, these spatial findings support mechano-inflammation, including a sustained stromal activation accompanied by an evolving macrophage response following local *Lats1/2* deletion.

### Local ablation of contractile machinery in synovial cells reduces hyperalgesia and synovial pathology after joint injury

Given the above findings indicating that synovial cell mechano-activation was sufficient to cause joint disease in the absence of injury, we next queried whether the opposite would also hold. That is, whether the elimination of key components of the contractile machinery would be sufficient to lessen disease progression following an instigating injury. Although we previously showed that intra-articular TAT-Cre efficiently recombined synovial cells in healthy knees, DMM induces a marked increase in local fibroblast and macrophage populations as well as the recruitment of circulating immune cells^38,39^, which could dilute or redistribute recombined cells during disease progression. To test the survival and persistence of recombined cells after injury, tdTomato Ai9 reporter mice received intra-articular PBS or TAT-Cre 3 and 1 days before DMM surgery. Flow cytometry at 1 week detected tdTomato in 16.3% of CD45^+^ synovial cells and 39.8% of CD45^−^ cells, compared to the 3.2% and 16.8% tdTomato^+^ cells detected within these gates in PBS-injected controls (**Supplementary Fig. 6A**). At 4 weeks, TAT-Cre-injected joints exhibited robust tdTomato fluorescence throughout the synovial lining and gutters, whereas PBS-injected joints showed minimal signal (**Supplementary Fig. 6B**). Thus, TAT-Cre-targeted stromal and immune populations persisted following DMM-induced joint injury.

We next used the TAT-Cre-mediated approach to determine whether reducing actomyosin contractility altered PTOA progression following injury. *Myh9* and *Myh10* encode the heavy chains of non-muscle myosin IIA and IIB, which generate cytoskeletal tension required for adhesion, migration, and mechanotransduction^11,40^. Our previous single cell data set identified significant increases in *Myh9/10* positive cell populations following DMM^21^, with similar increases observed in our Lats dKO mice in the absence of injury (**Supplementary Fig. 7**). To evaluate the role of this contractile machinery, *Myh9*^f/f^;*Myh10*^f/f^ mice received bilateral intra-articular PBS (hereafter, PBS control) or TAT-Cre (hereafter, Myosin dKO) prior to unilateral DMM surgery, with the contralateral intact knee serving as an internal control (**Fig. 5A**).

**Fig. 5:**
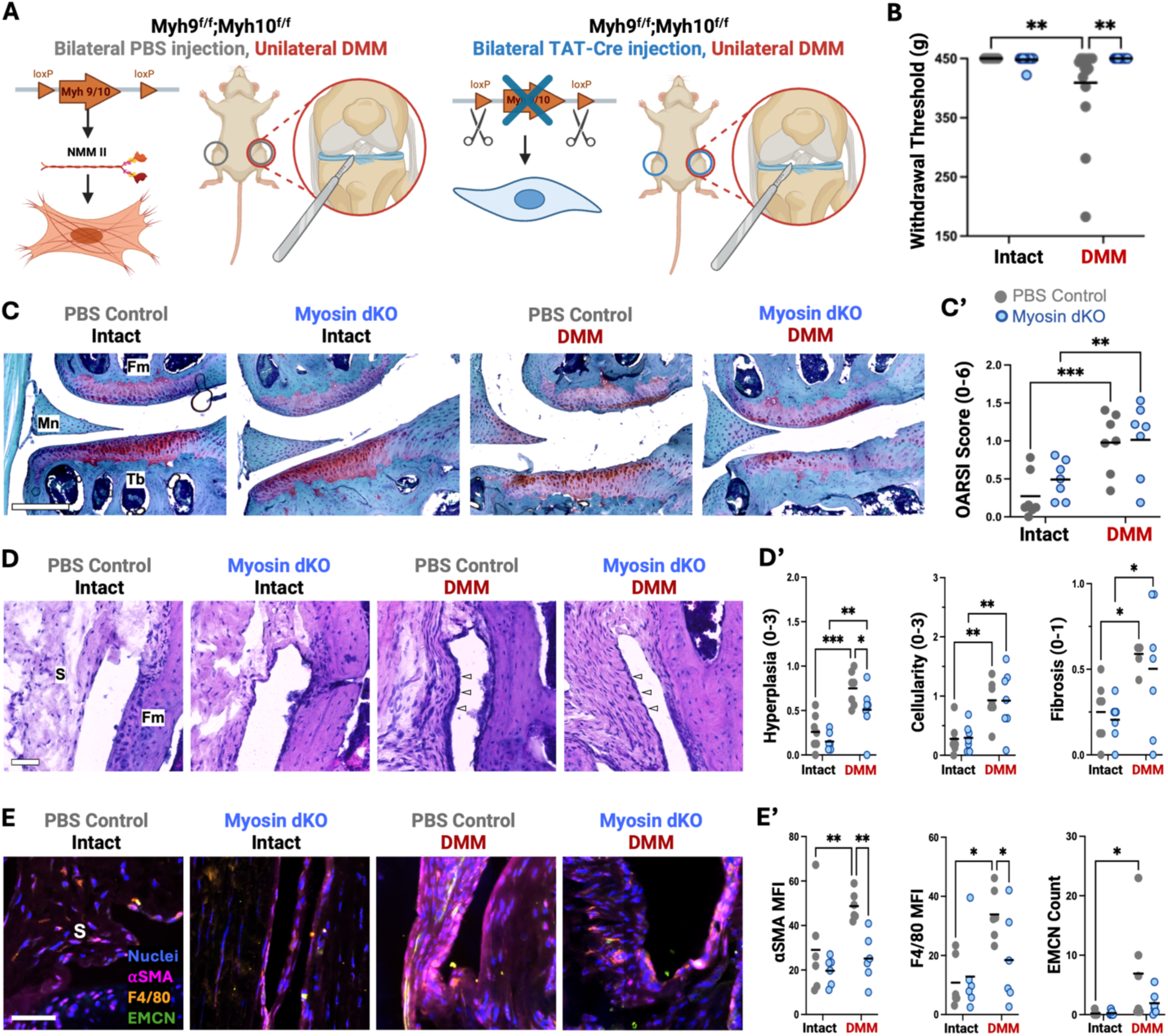
Local *Myh9/10* deletion reduces DMM-induced hyperalgesia and some features of synovial pathology. **(A)** Experimental design for bilateral intra-articular PBS or TAT-Cre administration in *Myh9/10* floxed mice followed by unilateral destabilization of the medial meniscus (DMM); contralateral knees served as intact controls. **(B)** Joint withdrawal thresholds in intact and DMM-operated knees 8 weeks after surgery. **(C)** Representative Safranin O/Fast green stained sections of articular cartilage (Scale = 250µm). **(C’)** OARSI scores for articular cartilage. **(D)** Representative H&E-stained synovial gutters. **(D’)** Semiquantitative scores for intimal hyperplasia, sublining cellularity, and fibrosis. **(E)** corresponding immunofluorescence images from PBS-intact, PBS-DMM, TAT-Cre-intact, and TAT-Cre-DMM knees. **(E’)** Immunofluorescence synovial lining quantification of αSMA, F4/80, and EMCN^+^ vessel number. Nuclei are in blue, αSMA is in magenta, F4/80 is in orange, and EMCN is in green. C, cartilage; Fm, femur; MCL, medial collateral ligament; Mn, meniscus; S, synovium; Tb, tibia. *p<0.05, **p<0.01, ***p<0.001.

Pain withdrawal thresholds did not differ among Myosin dKO groups at baseline or before surgery. At 8 weeks following DMM, PBS control mice exhibited lower withdrawal thresholds in operated knees than in contralateral intact knees (**Fig. 5B**), indicating increased hyperalgesia. In contrast, withdrawal thresholds did not differ between DMM-operated and intact knees in Myosin dKO mice and were higher in Myosin dKO DMM knees than in PBS control DMM knees. Thus, local *Myh9/10* deletion mitigated DMM-induced joint hyperalgesia.

Given the reduction in DMM-induced hyperalgesia, we next assessed whether this symptomatic improvement was accompanied by protection of the articular cartilage. OARSI scores increased after DMM in PBS control mice (**Fig. 5C, D**), as expected. Myosin dKO mice showed a similar numerical increase after DMM, and OARSI scores did not differ between PBS control and Myosin dKO DMM knees. Therefore, *Myh9/10* deletion did not detectably protect the cartilage at the 8-week endpoint.

We next examined the synovium to determine whether changes in this tissue accompanied the observed reduction in hyperalgesia. Histological scoring showed that DMM increased intimal lining hyperplasia, sublining cellularity, and fibrosis in both PBS control and Myosin dKO mice (**Fig. 5E, F**). However, lining hyperplasia was lower in Myosin dKO DMM knees compared to PBS control DMM knees. Sublining cellularity and fibrosis did not differ between treatment groups (**Fig. 5E**). Immunofluorescence further showed lower αSMA- and F4/80-associated fluorescence in Myosin dKO DMM joints compared to PBS control DMM joints (**Fig. 5E, F**). The number of EMCN^+^ vascular structures also showed a downward trend after *Myh9/10* deletion but did not differ significantly between treatment groups. These findings indicate that *Myh9/10* deletion reduced lining expansion, contractile stromal activation, and macrophage accumulation rather than broadly preventing all components of synovial remodeling.

Synovitis is closely associated with joint pain in osteoarthritis^31,33,41^. The reduction in hyperalgesia, together with lower lining hyperplasia, αSMA, and F4/80, despite unchanged cartilage scores, identifies a dissociation between symptomatic and structural outcomes in this model. Collectively, the *Lats1/2* gain-of-mechanosignaling and *Myh9/10* loss-of-contractility experiments support a causal role for synovial mechanobiology in causing joint pain and remodeling of the synovial cell landscape, while also showing that partial suppression of this pathway is insufficient to prevent cartilage degeneration over the interval examined in the context of a destabilizing injury.

## DISCUSSION

Here, we identify dysregulated synovial cell mechanotransduction as an active driver of joint disease, rather than a simple consequence of tissue damage. Local *Lats1/2* deletion was sufficient to induce sustained hyperalgesia, altered gait, and progressive synovial, vascular, and cartilage pathology in the absence of a destabilizing injury. Conversely, local *Myh9/10* deletion partially rescued PTOA pathogenesis, with reduced DMM-induced hyperalgesia, lining hyperplasia, contractile stromal remodeling, and macrophage accumulation. However, this intervention was unable to completely reverse pathologic changes in the joint, including continued cartilage deterioration. Together, these orthogonal perturbations demonstrate that increasing synovial mechanosignaling initiates joint-wide mechano-inflammatory pathology and contributes to joint degeneration. These findings extend prior work linking matrix stiffening and mechanoactivation to osteoarthritic synovium, in which mechanoactivated fibroblasts lose homeostatic functions such as lubricin production that can be restored in vitro by inhibiting Rho– ROCK-dependent contractility^23^, and in which disrupting myosin- or integrin-dependent mechanosignaling in vivo mitigates degeneration after joint injury^42–44^. However, most prior studies examined mechanosignaling after injury or targeted single downstream effectors^42–44^. By removing the upstream Hippo kinases *Lats1/2* in an otherwise uninjured joint, we isolated the loss of mechanosignaling restraint as an initiating event and showed that this was sufficient on its own to produce pain, functional impairment, and progressive multi-tissue pathology.

*Lats1/2* deletion did not act through a single effector population but reorganized the local stromal–immune cellular landscape. Deletion of *Lats1/2* in synovial cells increased mechanosensing and YAP/TAZ-associated transcriptional programs in both CD45^−^ and CD45^+^ compartments, coinciding with the emergence of *Cthrc1^+^* matrix-remodeling fibroblasts and *Trem2^+^Mrc1^+^*macrophages, populations linked to pathological remodeling in fibrotic tissue and arthritic pannus^35–37^, and to tissue repair^34^, respectively. Fibroblasts and tissue-resident macrophages are the dominant resident populations of the synovium and are increasingly recognized as mechanosensitive, adapting their phenotype and inflammatory output to changes in matrix stiffness^13–15^. The concurrent emergence of these two cell populations, alongside sustained fibroblast-associated remodeling and a more transient macrophage response, raises the possibility that their mechanosensitivity is coupled: reciprocal mechano-inflammatory cytokine signaling and matrix remodeling may link fibroblast and macrophage activation, such that mechanically induced changes in one population propagate pathological responses through the other. The present data cannot answer this question, given that TAT-Cre delivery recombines a fraction of cells across both compartments simultaneously and cannot separate cell-autonomous from secondary effects. These targeted approaches will be the focus of future work.

Comparing the *Lats1/2* and *Myh9/10* models shows that the site of pathway perturbation determines the outcome. *Lats1/2* normally restrains activity within broad YAP/TAZ-dependent network, and its loss permits coordinated shifts in cell state, matrix remodeling, and inflammatory signaling. *Myh9/10*, on the other hand, acts downstream of multiple adhesion and signaling inputs to generate actomyosin tension (**Fig. 6A**). Deletion of *Myh9/10* attenuated hyperalgesia, lining expansion, αSMA remodeling, and macrophage accumulation, but did not attenuate sublining fibrosis or prevent cartilage pathology (**Fig. 6B, C**). Thus, reducing cytoskeletal contractility appeared to ameliorate certain pathologic changes without suppressing the broader signals that sustain post-traumatic disease. This is consistent with the known dissociation between joint pain and structural degeneration^45^, and suggesting that synovial mechanotransduction may contribute more directly to nociceptive and inflammatory remodeling within the synovium rather than to cartilage preservation at this stage. Given the dramatic shifts in loading patterns following DMM (including higher stresses on the cartilage itself), restabilizing the joint in combination with inhibition of mechano-inflammation of the synovium may be required to curb PTOA progression.

**Fig. 6:**
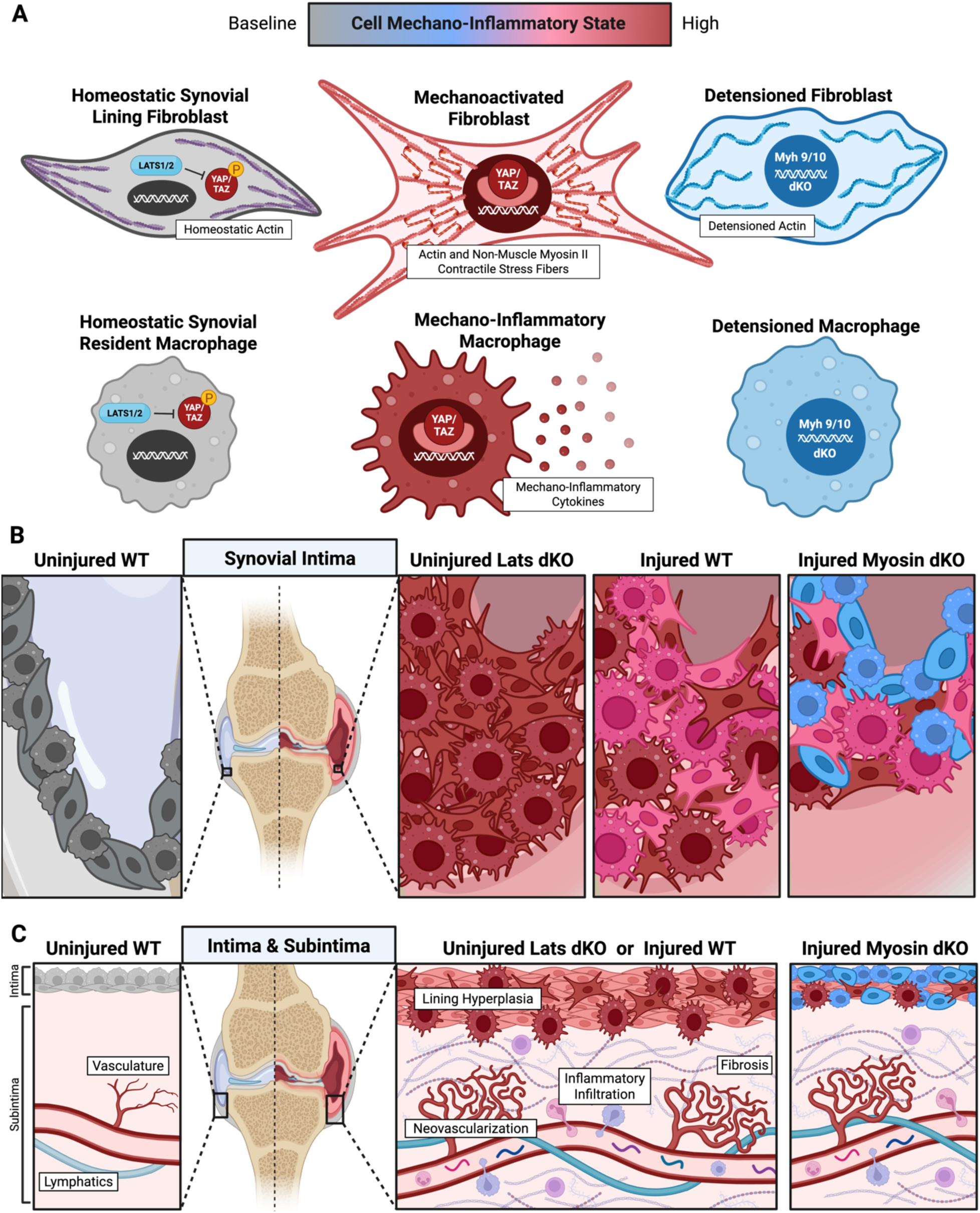
Actomyosin–YAP/TAZ mechanotransduction axis as a driver of synovial pathology. Schematic illustrating the proposed relationship between cellular mechanoactivation and synovial remodeling on a cellular and structural level. (**A**) Synovial fibroblast and macrophage phenotypes during baseline homeostasis (grey), a high mechano-inflammatory state (red), and a detensioned state (blue). During homeostasis, Lats1/2 restrain YAP/TAZ activity through phosphorylation and cytoplasmic retention or degradation. Joint injury or local *Lats1/2* deletion promotes mechanotransduction in both fibroblasts and macrophages characterized by actomyosin stress fibers, nuclear YAP/TAZ activity, and production of mechano-inflammatory cytokines. Local *Myh9/10* deletion prevents the cell from establishing stress fibers, thereby suppressing mechanotransduction. (**B**) Schematic of synovial lining cells, where *Lats1/2* deletion without injury causes the greatest increase mechano-inflammatory (red) cells. Injury in WT mice resulted in moderately increased mechano-inflammatory (red and pink) cells. *Myh9/10* deletion prior to injury results in detensioned cells (blue), fewer mechano-inflammatory cells, and decreased lining hyperplasia. (**C**) Schematic of synovial intima and subintima, showing increased neovascularization, inflammatory infiltration, and fibrosis in all experimental groups.

The two models also differ in timing: i) *Lats1/2* recombination preceded pathology, allowing a small, perturbed population to seed a feed-forward loop of activation and remodeling, ii) whereas *Myh9/10* deletion preceded DMM, after which proliferating or newly recruited cells outside the recombined pool could still sustain fibrosis and cartilage damage. This distinction matters therapeutically: releasing an upstream signaling brake tests whether mechanosignaling is sufficient to initiate disease, whereas inhibiting a downstream effector more closely models a partial intervention within an established injury response. Additionally, TAT-Cre targets only a small population of cells following injection that are sustained within the joint following injury. It may be that a small population of aberrant cells is sufficient to propagate fibrotic and inflammatory signaling in adjacent non-recombined cells, whereas the reverse is not true with ablation of contractility in only a small population of recombined cells in the otherwise larger mechano-activated population that emerges following injury.

These findings raise three important questions what will guide future studies. First, macrophage infiltration peaked early and declined by 4 WPI following Lats depletion, whereas αSMA-associated remodeling remained elevated throughout. This suggests that macrophages may help initiate the phenotype while contractile stromal populations sustain it, although parallel, cell-autonomous responses in both compartments remain equally plausible. Resolving this will require cell-restricted *Lats1/2* and *Myh9/10* perturbation in fibroblast and myeloid lineages, together with temporally controlled depletion and co-culture studies, to define which population initiates the response, whether either is necessary for activating the other, and more broadly, how cytoskeletal mechanosensing is coordinated across stromal and immune compartments to drive pathological remodeling. The second important question whether the *Cthrc1^+^* fibroblast and *Mrc1^+^ Trem2^+^* macrophage states are pathogenic, compensatory, or simply stage-dependent remains to be tested directly. Third, the severity of the *Lats1/2* phenotype suggests that joint homeostasis depends on maintaining mechanosignaling within a bounded range. Pharmacological studies will be essential to test this hypothesis, since neither genetic model addresses the treatment of established disease. Temporally staged inhibition of Rho–ROCK, actomyosin, and/or YAP/TAZ– TEAD signaling, initiated after pathology is established, would test whether remodeling can be reversed and whether targeting fibroblasts, macrophages, or both is required to preserve joint structure while relieving pain. Together, these findings position mechanotransduction as an intrinsic component of joint disease biology and provide a framework for identifying the cellular and pathway-specific interventions needed to uncouple maladaptive synovial remodeling from normal mechanical homeostasis.

## METHODS

### Animals and study design

All animal experiments were approved by the University of Pennsylvania Institutional Animal Care and Use Committee (protocol 806669). Prg4Cre^ERT^^2^;R26R^tdTomato^ (Ai9) reporter mice, Lats*1*^f/f^;*Lats2*^f/f^ mice and wild-type C57BL/6 mice were used to assess intra-articular recombination and the effects of local *Lats1/2* deletion. *Myh9*^f/f^;*Myh10*^f/f^ mice were used to assess the effects of local non-muscle myosin II deletion in destabilization of the medial meniscus (DMM)-induced osteoarthritis. The *Lats1/2* and *Myh9/10* conditional lines lacked a constitutive Cre driver; local recombination was induced by intra-articular TAT-Cre delivery. Male and female 12-week-old mice were used for all experiments.

For reporter validation in uninjured joints, tdTomato Ai9 mice received bilateral PBS or TAT-Cre injections and were euthanized 1 week after the second injection. seven mice were used per treatment, with three used for fluorescence imaging and four for flow cytometry.

For local *Lats1/2* deletion, one knee received TAT-Cre and the contralateral knee received PBS. Tissue endpoints were assessed at 1, 2 and 4 weeks after injection. Wild-type mice received the same unilateral treatment and contralateral PBS control for behavioral comparisons. Gait was assessed before treatment and at 1, 2 and 4 weeks after second TAT-Cre injection.

For DMM experiments, tdTomato Ai9 and *Myh9*^f/f^; *Myh10*^f/f^ mice received PBS or TAT-Cre 3 and 1 days before surgery. In the *Myh9/10* experiment, both knees received the assigned injection treatment, DMM was performed on the right knee and the contralateral knee remained intact. Reporter fluorescence was assessed 4 weeks after DMM. Reporter flow cytometry was assessed at 1 week post DMM. Pain (n=16), histopathology (n=7), and immunofluorescence (n=6) in the *Myh9/10* cohort were assessed 8 weeks after DMM.

### Intra-articular TAT-Cre delivery

TAT-Cre (Excellgen, EG-1001) was administered in two intra-articular injections 48 h apart. Each knee received 10 µl of TAT-Cre at 120 µM or an equal volume of sterile PBS, using a 25-µl Hamilton syringe with a 33-gauge needle (65460-11).

### Destabilization of the medial meniscus

DMM was performed on the right knee as previously described. Mice were maintained under 3% isoflurane anesthesia delivered in oxygen at 2-3 L min⁻¹, and Ethiqa XR was administered subcutaneously prior to surgery at 3.25 mg/kg for analgesia. Under a dissecting microscope, an incision was made along the medial border of the patellar tendon, and the joint capsule was opened. The infrapatellar fat pad was bluntly dissected to expose the medial meniscotibial ligament, which was transected with a no. 11 blade. Destabilization was confirmed by medial displacement of the meniscus. The capsule and skin were closed separately with 7-0 Vicryl sutures (Med-Vet International, J488G).

### Knee pressure sensitivity

Knee withdrawal thresholds were measured using a pressure application measurement device (Ugo Basile), as previously described. Pressure was applied to the knee at 30 g/s, with a maximum of 450 g. The pressure at limb withdrawal was recorded. The procedure was performed twice per knee. *Lats1/2* and wild-type cohorts were assessed at baseline, 1, 2 and 4 weeks after injection. The *Myh9/10* DMM cohort was assessed at baseline, before surgery and 8 weeks after surgery.

### Gait analysis

Gait was assessed in *Lats1*^f/f^; *Lats2*^f/f^ and wild-type C57BL/6 mice using the CatWalk XT system (version 10.6 Nodulus Information Technology, Wageningen, Netherlands) 1 week before injection and at 1, 2 and 4 weeks after TAT-Cre injection, as previously described. Mice acclimated to the testing environment for 30 min before data collection. Valid runs consisted of uninterrupted forward movement across the 1.3-m glass walkway without stopping, turning or reversing direction. Three to five valid runs were included per mouse at each time point. Automated identification of the four paws was manually verified. Recorded parameters included paw contact area, print width, stand time, swing time, swing speed and duty cycle, together with run duration, speed and cadence. Principal component analysis was used to summarize gait variation across genotypes and time points.

### In vitro reporter fluorescence imaging

Primary synovial fibroblasts were harvested from the hind paws of 12-week-old Prg4Cre^ERT^^2^;R26R^tdTomato^ (Ai9) reporter mice (n = 3) as previously described in *Knights et al*.^46^ Briefly, hind paws were prepared by removing the skin, claws, muscle, and tendon. Longitudinal cuts were then made along the hand paw using a #11 blade. Dissected hind paws were digested for 45 minutes at 37°C in 2mL of digestion media comprising DMEM, 400μg/mL collagenase IV (Worthington Biochemical Corp, LS004188), 400μg/mL Liberase TM Research Grade (Roche, 05401127001), and 400μg/mL DNase I (Roche, C756Z68), vortexing every 5-10 minutes. Following digestion, the cell suspension was filtered through a 40μm strainer and centrifuged at 500xg for 5 minutes. The resulting pellet was resuspended in basal media (DMEM with 10% fetal bovine serum and 1% penicillin/streptomycin/fungizone) and plated on tissue culture polystyrene.

Cells harvested from tdTomato Ai9 mice were plated on glass coverslips in 24-well plates (3.5k cells/well) at passage 3 in basal media (n = 3). After 24 hours, cells were treated with TAT-Cre (3μM, Excellgen EG-1001) for 5 hours. The media was then removed, and cells were rinsed twice with phosphate-buffered saline (PBS) before fresh media was added. Forty-eight hours after TAT-Cre treatment, cells were fixed in microtubule stabilizing buffer (2% paraformaldehyde, 0.1M PIPES (pH 6.75), 1mM EGTA, 1mM MgSO4, 4% (w/v) poly(ethylene glycol), 1% Triton X-100) and coverslips were mounted with ProLong Gold Antifade Mountant with DAPI (ThermoFisher, P36931). Cells were imaged at 20x on a Zeiss Axio Scan.Z1 Slide Scanner (Zeiss, Oberkochen, Germany). The percentage of cells that were tdTomato-positive was calculated in 4 representative regions per coverslip and averaged.

### Tissue processing and reporter whole-joint fluorescence imaging

Harvested hindlimbs were fixed in 10% neutral buffered formalin for 24 h. TdTomato Ai9 reporter joints were decalcified in 14% EDTA p.H. 7.4 for 72h; all other joints were decalcified in Formical for 7 days. Next, joints were incubated in 30% sucrose for 48 h and embedded in optimal cutting temperature compound. Coronal or sagittal, tape-stabilized cryosections were collected at 8 µm thickness. Sections on cryofilm were attached to glass slides using 0.75% chitosan (Sigma-Aldrich, 419419) in 0.25% acetic acid (Sigma-Aldrich, 695092) and dried overnight at room temperature.

For reporter whole-joint imaging, coronal and sagittal sections were mounted with Hoechst 33342 in 50% glycerol (Invitrogen, 15514011) 50% PBS, and imaged with a ×20 objective on a Zeiss Axio Scan.Z1 slide scanner. Native tdTomato fluorescence was used to visualize recombination in Ai9 joints.

### Reporter flow cytometry

One week after TAT-Cre or PBS injections, in joints with or without DMM, synovial tissue was dissected from four tdTomato Ai9 mice and pooled to generate 1 sample per condition. Synovium was digested for 35 minutes at 37°C in DMEM containing liberase (1 U ml−1) and DNase I (0.18 mg/mL). Cell suspensions were passed through a 40-µm strainer and centrifuged at 500 × g for 5 min. Pellets were resuspended in PBS containing 2% bovine serum albumin and 1 mM EDTA. Red blood cells were lysed using RBC lysis buffer (Invitrogen, 00-4333-57), and viability was assessed using LIVE/DEAD Fixable Aqua stain (Thermo Fisher Scientific, L34957), according to the manufacturers’ instructions.

Cells were incubated with Alexa Fluor 488 anti-mouse CD45 antibody (BioLegend 103121, raised in rat, diluted 1:100) for 30 min on ice in the dark, washed and resuspended for acquisition.

Controls included unstained cells, unstained tdTomato-positive fibroblasts, viability-stained dead cells, and positive and negative compensation beads. Compensation controls were first acquired, and then synovial data were acquired on a BD LSRFortessa Flow Cytometer (BD Biosciences, Franklin Lakes, New Jersey) and analyzed in FlowJo v10.9.0 (BD/Treestar). Cells were gated to exclude debris, doublets, and dead cells. Then, gated according to CD45 and tdTomato fluorescence. Recombination was assessed as the percentage of tdTomato-positive cells within the designated CD45 gate.

### Histological staining and scoring

*Lats1/2* joints were assessed at 1, 2 and 4 weeks after injection, and *Myh9/10* joints were assessed 8 weeks after DMM. Cryosections were stained with hematoxylin and eosin to assess joint and synovial morphology. Synovial lining hyperplasia, sublining cellularity and fibrosis were scored by four independent, blinded reviewers using a validated murine scoring system^47^. Scores were assigned in the medial femoral, medial tibial, lateral femoral and lateral tibial synovial gutters and averaged to obtain medial, lateral and whole-joint values.

Cartilage was stained with Safranin O/Fast Green and scored from 0 to 6 by four independent, blinded reviewers in each quadrant according to the OARSI mouse histopathology guidelines^48^. Medial, lateral and whole-joint averages were calculated. *Lats1/2* joints were also stained with Toluidine Blue to identify glycosaminoglycans.

### Lats 1/2 immunofluorescence and image quantification

Coronal joint cryosections (8 µm) underwent permeabilization with PBS containing 0.4% Triton X for 15 min at room temperature and were blocked with 5% bovine serum albumin diluted into PBS containing 0.1% Triton X for 60 min at room temperature. Samples were incubated with F4/80 and Endomucin primary antibodies overnight at 4 °C (Table 1). After three washes with PBS, samples were incubated with anti-rat and anti-goat secondary antibodies for 60 min at room temperature. After three washes with PBS, samples were incubated with anti-αSMA antibody and Hoechst for 60 min at room temperature (Table 1). After three washes with PBS containing 0.1% Triton X, samples were incubated in PBS overnight, mounted with 50% glycerol 50% PBS, and imaged with a ×20 objective on a Zeiss Axio Scan.Z1 slide scanner.

**Table 1.**
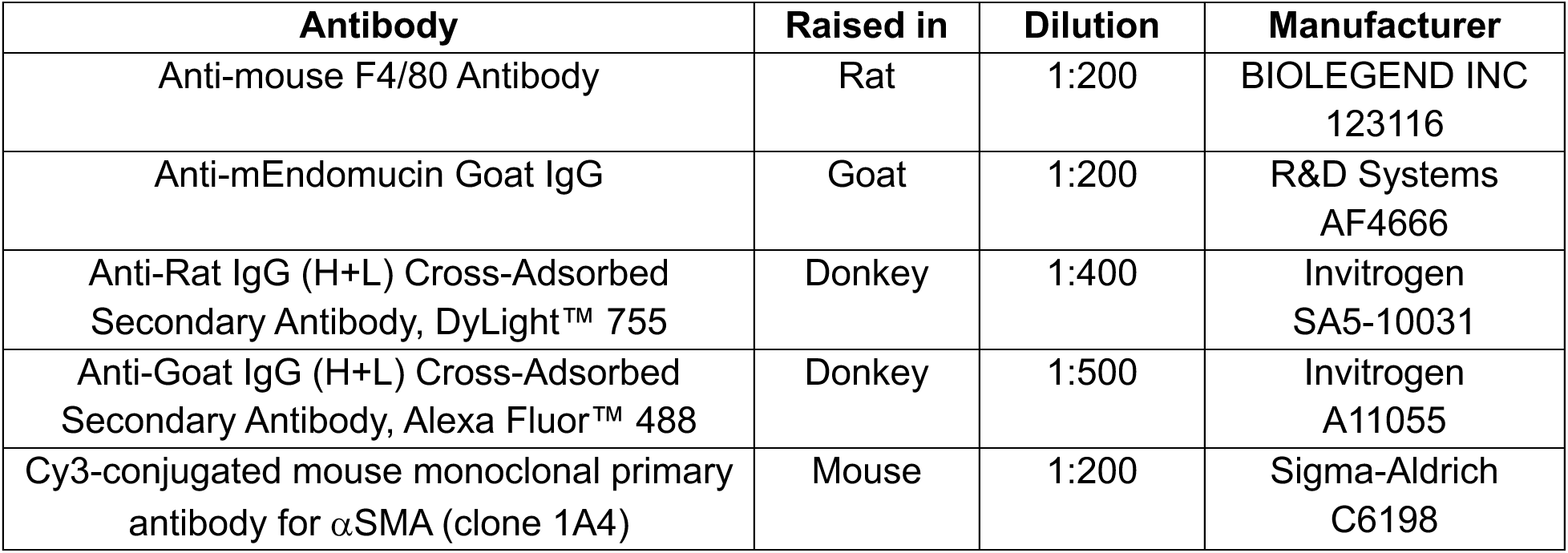
*Lats1/2* IF Antibodies.

Image analysis was performed in ImageJ. First, ROIs were manually drawn using the nuclear channel to exclude the bone, menisci, ACL and PCL, and to include the pathologic tissue in between the bone and muscle. Within this ROI, the mean fluorescence intensity (MFI) of F4/80 and αSMA channels was quantified. Background MFI was measured for the F4/80 and αSMA channels in each sample and averaged across samples. This background value was then subtracted from F4/80 and αSMA MFI for each sample. Within this ROI, the endomucin channel was thresholded and particles with area <10µm were excluded. Then, total number of endomucin-positive particles were automatically counted, and the percent endomucin-positive area was quantified.

### *Myh9/10* immunofluorescence and image quantification

Coronal joint cryosections (8 µm) underwent antigen retrieval with Proteinase K (Dako, S3020) for 5 min at room temperature and were blocked with Protein Block (Abcam, ab64226). Sections were incubated overnight at 4 °C with rat anti-F4/80, mouse anti-α-smooth muscle actin, and goat anti-endomucin (Table 2). After two washes with Tris-buffered saline, sections were incubated for 1 h at room temperature with donkey anti-goat Alexa Fluor 488 secondary antibody. Sections were washed twice and incubated for 1 h at room temperature with goat anti-mouse Alexa Fluor 546 and goat anti-rat Cy5 secondary antibodies (Table 2). After washing, sections were counterstained with Hoechst 33342, mounted with 50% glycerol 50% PBS, and imaged with a ×20 objective on a Zeiss Axio Scan.Z1 slide scanner.

**Table 2.** *Myh9/10* IF Antibodies.

| Antibody | Raised in | Dilution | Manufacturer |
| --- | --- | --- | --- |
| Monoclonal primary antibody for F4/80 | Rat | 1:200 | BioLegend<br>123102 |
| Monoclonal primary antibody for αSMA (clone 1A4) | Mouse | 1:200 | Sigma-Aldrich<br>A2547 |
| Polyclonal primary antibody for endomucin | Goat | 1:100 | R&D Systems<br>AF4666 |
| Anti-goat Alexa Fluor 488 | Donkey | 1:500 | Invitrogen<br>A11055 |
| Anti-mouse Alexa Fluor 546 | Goat | 1:500 | Invitrogen<br>A11030 |
| Anti-rat Cy5 | Goat | 1:500 | Invitrogen<br>A10525 |

F4/80 and α-smooth muscle actin fluorescence intensity was quantified in ImageJ within each of the four synovial gutters and averaged across the joint. Endomucin-positive vessels were counted manually in the same regions and averaged across the joint.

### Single-cell RNA sequencing

Whole knee joints were collected 2 weeks after PBS or TAT-Cre injection. Twelve animals were used per treatment, with six males and six females pooled to generate one sample per condition.

Excised joints were placed in DMEM containing 1% penicillin/streptomycin/fungizone, and most of the surrounding muscle was removed. The patella and associated tendon were removed. The distal femur and proximal tibia were transected, and bone marrow was flushed. Remaining joint tissues (including the synovium, menisci, collateral ligaments, articular cartilage, fat pad, muscle, and ends of the femur and tibia) were minced and centrifuged at 500 × g for 5 min. Tissue was resuspended in DMEM containing 1% penicillin/streptomycin/fungizone, collagenase P (2 mg ml−1, Sigma Aldrich, 11249002001), hyaluronidase (2 mg ml^−1^, Sigma Aldrich, H3506) and DNase I (0.18 mg ml^−1^, Sigma Aldrich, DN25), and digested for 30 min in a shaking water bath at 37 °C with intermittent mixing.

Following digestion, samples were diluted in PBS containing 0.04% bovine serum albumin, dissociated by repeated pipetting and sequentially filtered through 70- and 40-µm strainers. Cells were centrifuged at 400 × g for 10 min and incubated in red blood cell lysis buffer for 2 min at room temperature. Lysis was quenched with cold PBS containing 0.04% bovine serum albumin, and cells were centrifuged at 400 × g for 10 min. Dead cells were removed using Dead Cell Removal MicroBeads (Miltenyi Biotec) and magnetic columns according to the manufacturer’s instructions. Live cells were washed and resuspended in PBS containing 0.04% bovine serum albumin. Cell concentration and viability were assessed by trypan blue exclusion.

Cell suspensions with viability exceeding 90% were processed by the Children’s Hospital of Philadelphia Center for Applied Genomics using the Chromium Next GEM Single Cell 3′ Kit v3.1 (10x Genomics). Libraries were indexed with the Chromium Dual Index Kit, pooled and sequenced using paired-end, dual-index sequencing on an Illumina NovaSeq. Approximately 20,000 reads per cell were generated. Demultiplexing, alignment to the mouse mm10 reference and feature–barcode matrix generation were performed using Cell Ranger v6.1.2.

### Single-cell data analysis

Feature–barcode matrices were analyzed using Seurat in R. Cells with fewer than 500 detected genes or fewer than 500 unique molecular identifiers (UMIs) were excluded. Additional filters excluded cells exceeding, a mitochondrial fraction above 5% or a hemoglobin fraction above 1%. Cells within the lowest 8% of detected-gene counts or UMI counts were also excluded.

Data were log-normalized using NormalizeData. Principal components were selected using elbow plots and principal-component heatmaps. Nearest-neighbor graphs were constructed using FindNeighbors with dimensions 1–34, and clusters were identified using FindClusters at resolution 0.1. Uniform manifold approximation and projection (UMAP) was used for visualization.

Cluster markers were identified using FindAllMarkers. The source specifies selection of genes expressed in more than 70% of cells within a cluster and fewer than 30% of cells outside that cluster. Cluster identities were assigned using canonical markers and inspected using violin and feature plots.

Cells were divided into immune and non-immune compartments using *Ptprc* expression greater than 1 to define the immune subset. Each subset was independently normalized, subjected to variable-feature selection and scaled before PCA. FindNeighbors used dimensions 1–25; FindClusters used resolutions of 0.1 and 0.07 for the immune and non-immune subsets, respectively. UMAP was used for visualization. Cluster proportions were calculated within each treatment and compartment. YAP/TAZ-responsive gene expression was visualized in heatmaps for the immune and non-immune compartments.

### Statistical analysis

Statistical analyses were performed in GraphPad Prism v10.3.0. The source draft specifies outlier assessment using the ROUT method (Q = 1%) and normality assessment using the Shapiro–Wilk test. Paired comparisons between PBS- and TAT-Cre-injected knees in the *Lats1/2* cohorts were analyzed using paired t-tests for normally distributed data and Wilcoxon matched-pairs signed-rank tests otherwise. Pain and gait comparisons across injection groups and genotypes were analyzed using repeated-measures two-way analysis of variance with Fisher’s least significant difference post hoc tests. *Myh9/10* DMM outcomes were analyzed using repeated-measures two-way analysis of variance with Fisher’s least significant difference post hoc tests.

## ACKNOWLEDGEMENTS

### Author contributions

Conceptualization: ERB, GK, JDB, NAD, RLM. Methodology: ERB, GK. Investigation: ERB, GK, AAS, TB, BC, RF. Formal analysis: ERB, GK, AAS. Visualization: ERB, GK, AAS. Writing – original draft: ERB, GK, AAS. Writing – review and editing: GK, ERB, AAS, LM, CS, JDB, NAD, RLM. Supervision: JDB, NAD, RLM. Funding acquisition: LM, CS, JDB, NAD, RLM. ERB and GK contributed equally to this work.

## SUPPLEMENTAL FIGURES and LEGENDS

**Supplementary Fig. 1:**
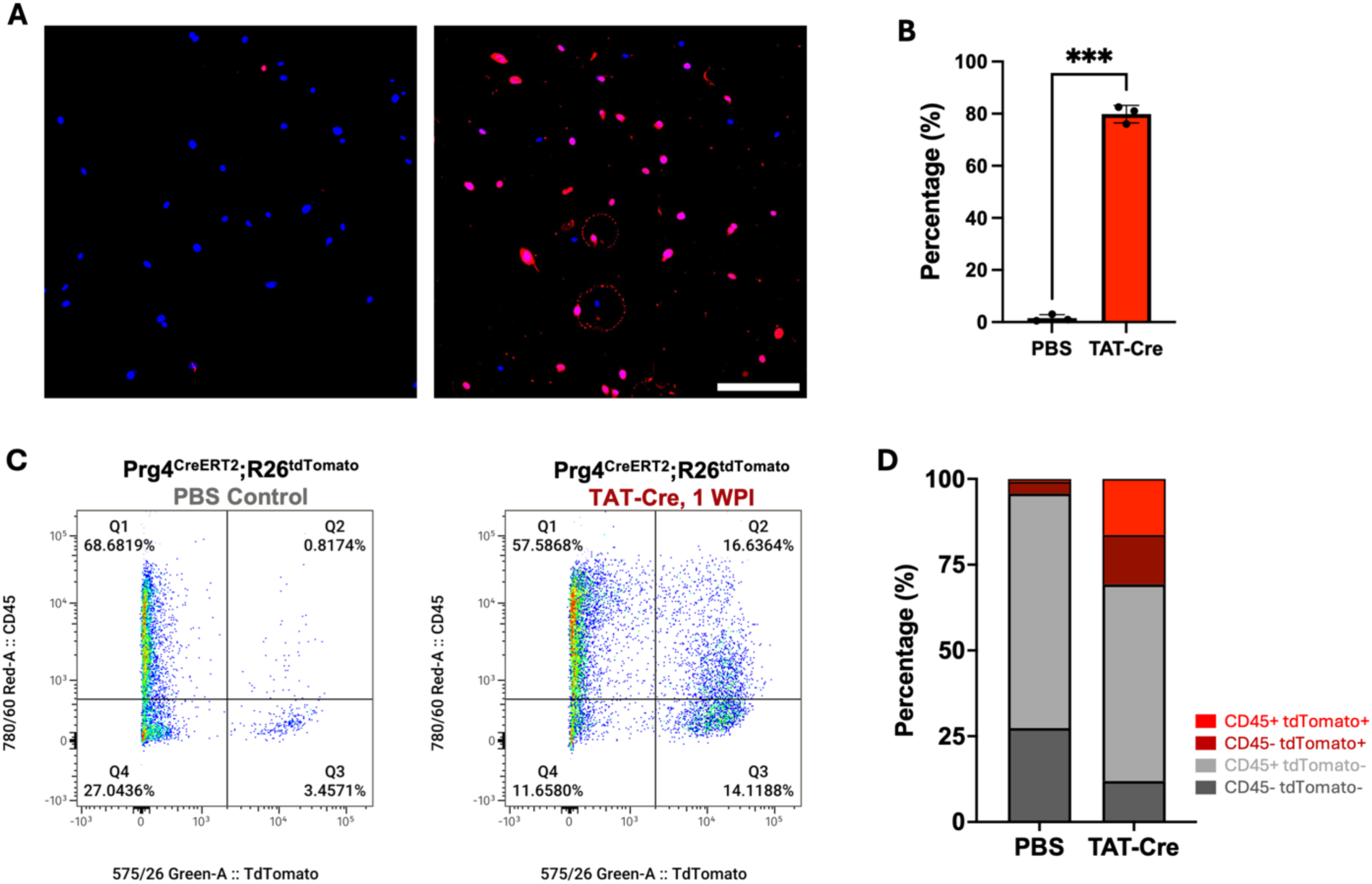
TAT-Cre validation in tdTomato Ai9 reporter mice. **(A)** Representative images of primary fibroblasts untreated or treated with TAT-Cre. Scale: 50μm. **(B)** Percentage of cells in each treatment group expressing tdTomato. ***p<0.001. **(C)** Flow cytometry plots and **(D)** graphical representation of synovial cells isolated from mice one week post injection with PBS or TAT-Cre, gated for CD45 and tdTomato expression.

**Supplementary Fig. 2:**
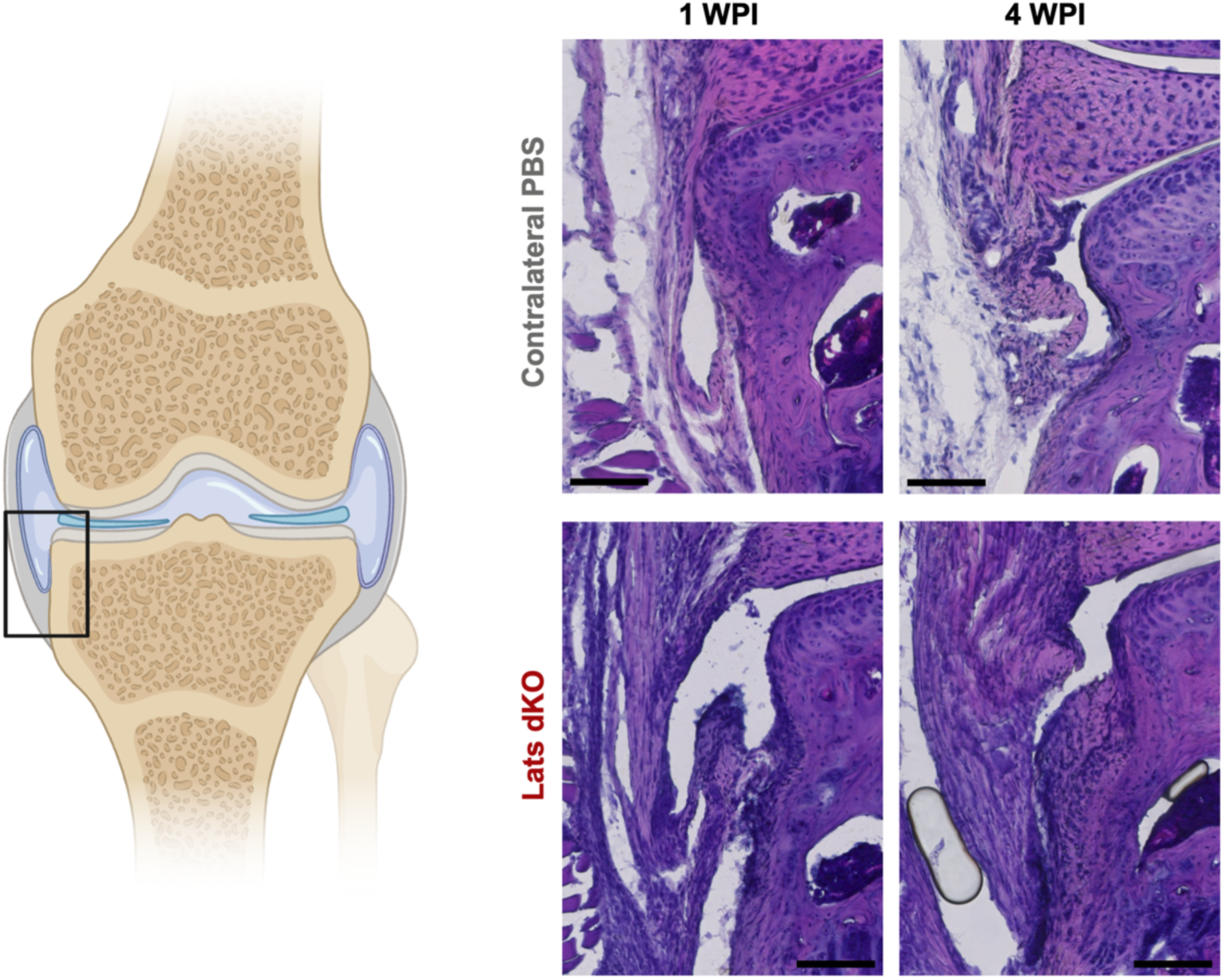
Lats dKO synovium. Additional representative H&E images of PBS or TAT-Cre injected Lats1/2 knees at 1 and 4 WPI. Scale = 100 μm.

**Supplementary Fig. 3:**
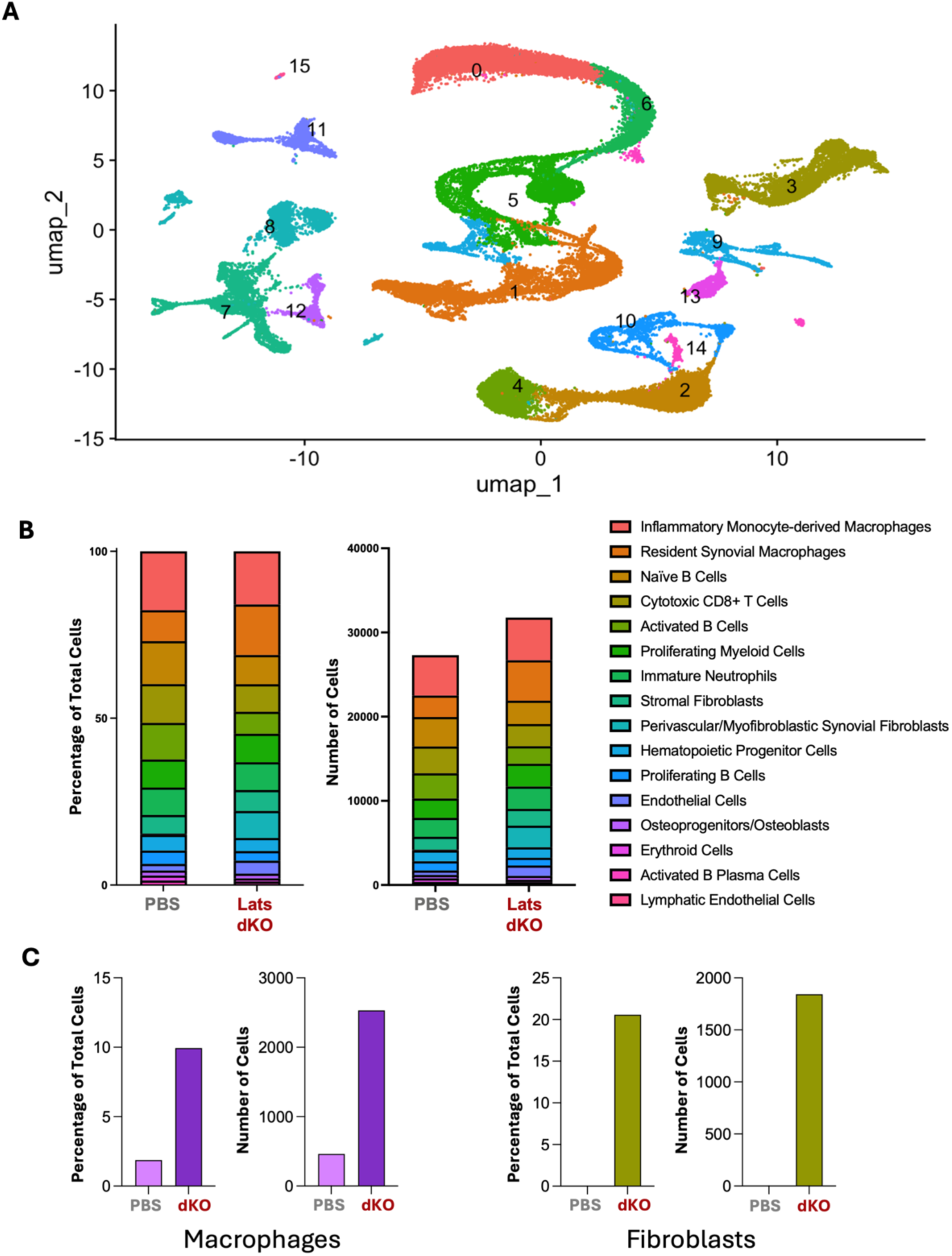
Single cell analysis of all cells isolated from PBS-injected control and TAT-Cre-injected Lats1/2 knees. **(A)** UMAP projection of all cells. Bar charts showing **(B)** subpopulation composition and **(C)** macrophage and fibroblast composition of PBS- and TAT-Cre-injected knees.

**Supplementary Fig. 4:**
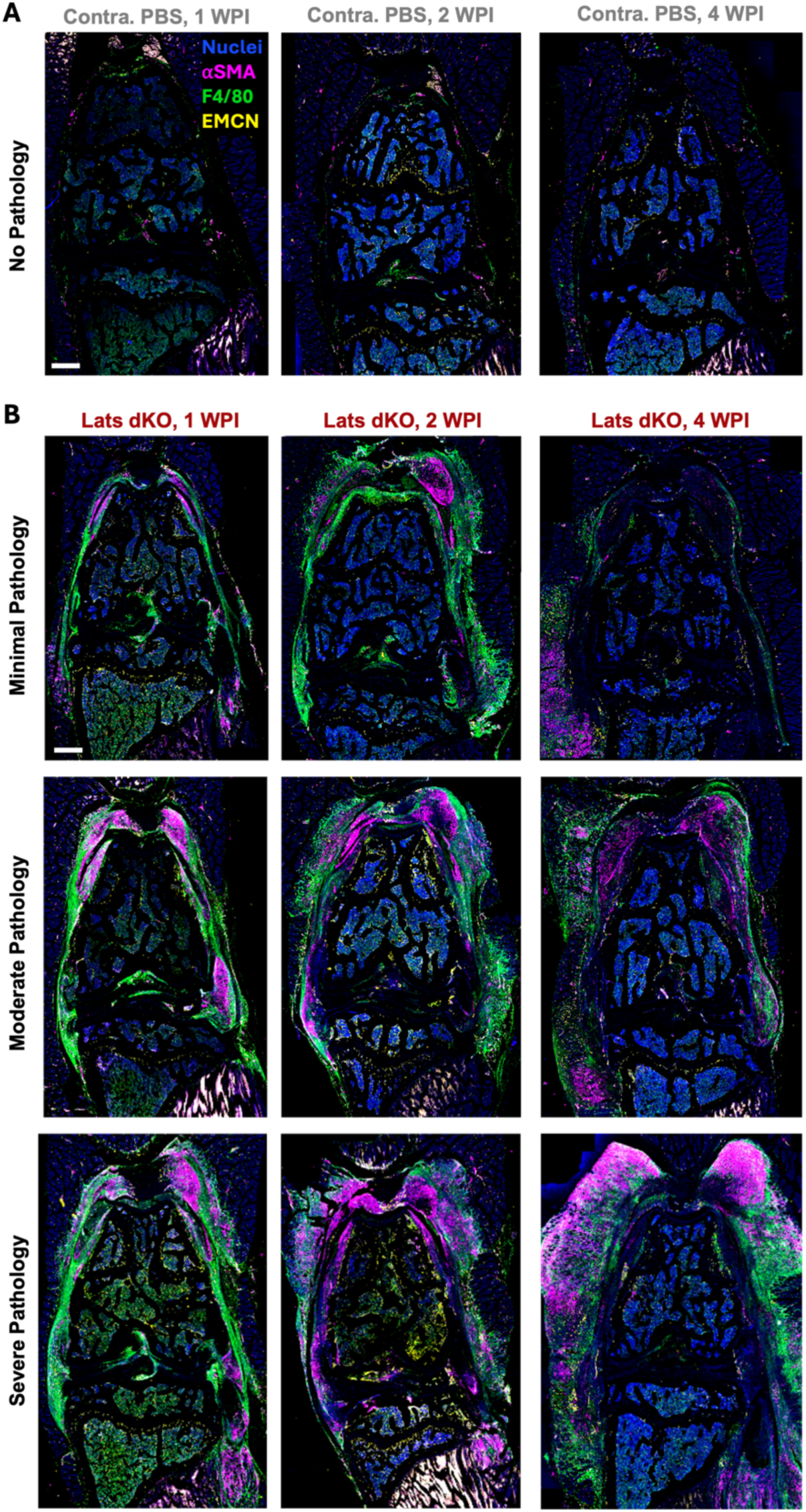
Representative immunofluorescence images showing varying degrees of Lats dKO pathology. **(A)** PBS-injected contralateral knees and **(B)** TAT-Cre-injected Lats1/2 knees at 1, 2 and 4 WPI. Nuclei are in blue, αSMA is in magenta, F4/80 is in green, and endomucin is in yellow. Scale = 500 μm.

**Supplementary Fig. 5:**
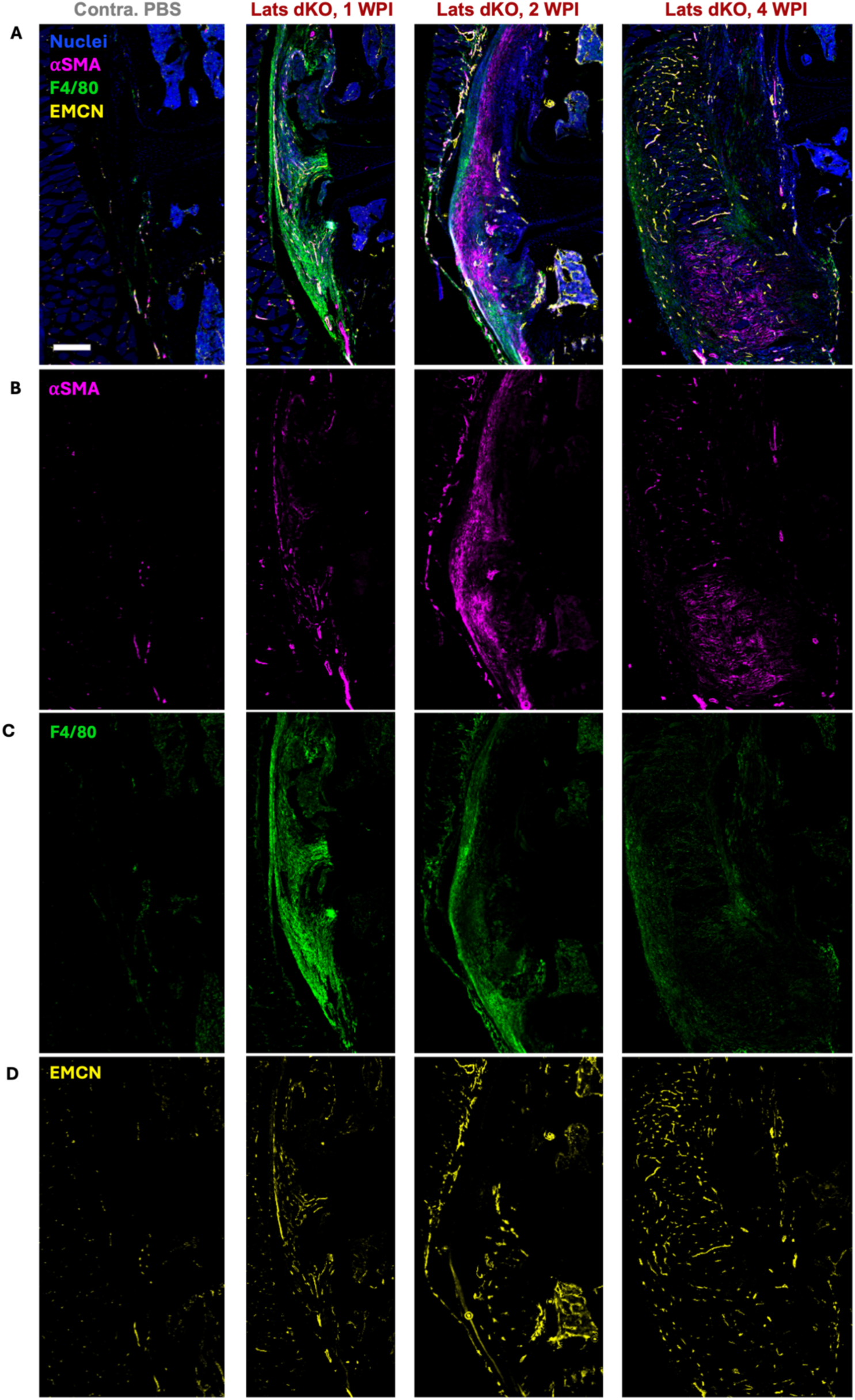
Representative immunofluorescence images of PBS-injected contralateral knees and TAT-Cre-injected Lats1/2 knees at 1, 2 and 4 WPI. Each row corresponds to a different immunofluorescence channel, **(A)** including all channels, **(B)** magenta αSMA, **(C)** green F4/80, and **(D)** yellow endomucin. Scale = 500 μm.

**Supplementary Fig. 6:**
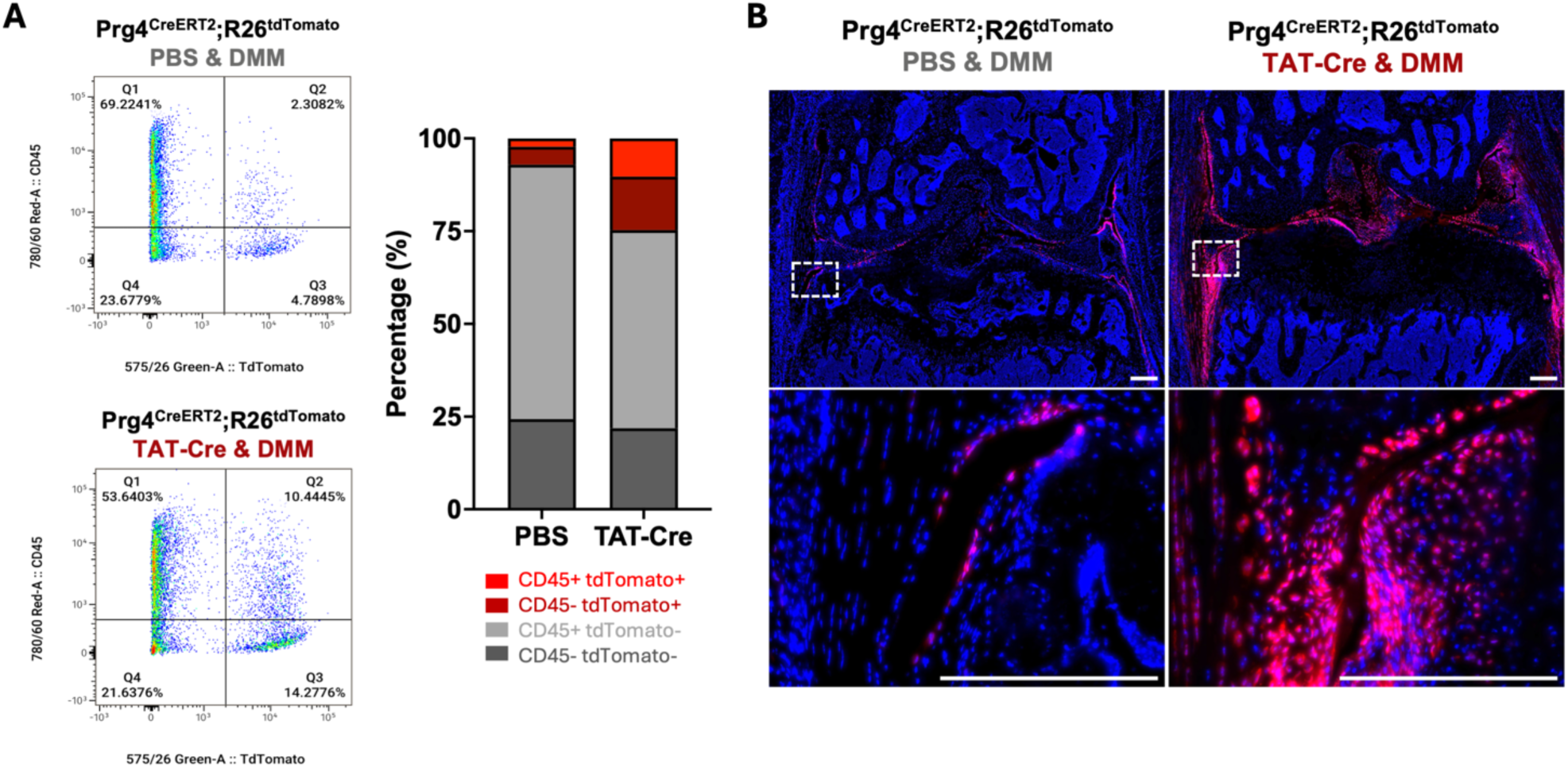
Persistence of TAT-Cre recombined cells following DMM. TdTomato Ai9 reporter mice were injected with PBS or TAT-Cre and then underwent DMM to produce OA. **(A)** Flow cytometry plots and graphical representation of synovial cells isolated from mice one week post DMM, gated for CD45 and tdTomato expression. **(B)** Immunofluorescence at 4 weeks post DMM, with the synovium highlighted by the dashed box (with zoom below). Nuclei are in blue and tdTomato reporter is in red. Scale = 250 μm.

**Supplementary Fig. 7:**
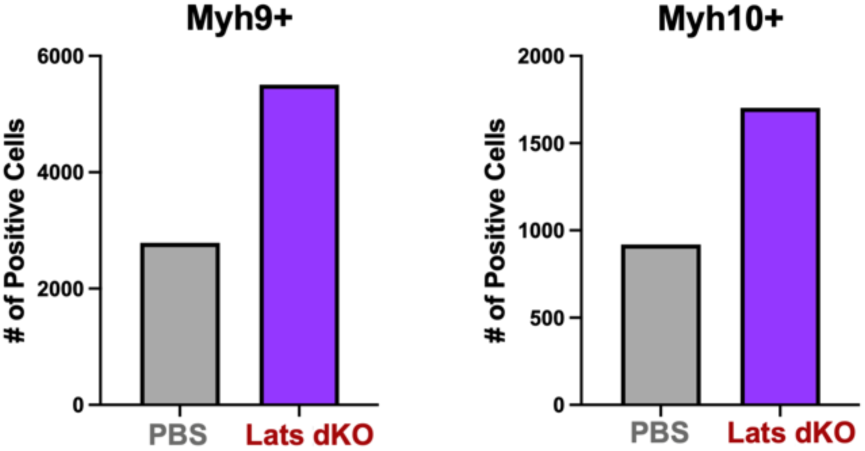
Myh9/10 expression in Lats1/2 single cell sequencing. Graphs show number of Myh9^+^ or Myh10^+^ cells identified in PBS or TAT-Cre injected Lats1/2 knees.

